# Downregulation of the silent potassium channel Kv8.1 increases ALS motor neuron vulnerability

**DOI:** 10.1101/2023.05.07.539778

**Authors:** Xuan Huang, Seungkyu Lee, Riki Kawaguchi, Ole Wiskow, Sulagna Ghosh, Devlin Frost, Laura Perrault, Roshan Pandey, Kuchuan Chen, Joseph R. Klim, Bruno Boivin, Crystal Hermawan, Kenneth J. Livak, Dan Geschwind, Brian J. Wainger, Kevin Eggan, Bruce P. Bean, Clifford J Woolf

**Affiliations:** F.M. Kirby Neurobiology Research Center, Boston Children’s Hospital, Boston, United States; Department of Psychiatry and Semel Institute for Neuroscience and Human Behavior, David Geffen School of Medicine, University of California Los Angeles, Los Angeles, United States; Department of Stem Cell and Regenerative Biology, Department of Molecular and Cellular Biology, and Harvard Stem Cell Institute, Cambridge, United States; Translational Immunogenomics Lab, Dana-Farber Cancer Institute, Boston, United States; Department of Neurology, Sean M. Healey & AMG Center for ALS, Massachusetts General Hospital, Harvard Medical School, Boston, United States; Department of Neurobiology, Harvard Medical School, Boston, United States

## Abstract

The Kv8.1 potassium ion channel encoded by the KCNV1 gene is a “silent” subunit whose biological function is unknown. In ALS patient-derived motor neurons carrying SOD1(A4V) and C9orf72 mutations, its expression is highly reduced, yielding increased vulnerability to cell death without a change in motor neuronal firing. Our data suggests that Kv8.1 modulates Kv2 channel function to impact neuronal metabolism and lipid/protein transport pathways, but not excitability.

## Main

Amyotrophic lateral sclerosis (ALS) is a devastating neurodegenerative disease that results in a rapid and progressive loss of motor neurons (MN) in the motor cortex and spinal cord ^1^. Advances in induced pluripotent stem cell (iPSC) technology not only enable in vitro modeling of ALS, but also facilitate the discovery of novel molecular targets in neurons ^2^. MNs differentiated from iPSCs derived from patients with sporadic and familial forms of ALS recapitulate key hallmarks of the disease, including increased cell death, shorter neurite outgrowth, protein aggregation, ER and unfolded protein stress responses, and abnormal excitability ^3-6^. Previously, our lab found that a positive feed-forward cycle of ER stress and abnormal excitability drives neuronal death in patient-derived motor neurons harboring the familial ALS A4V mutation in superoxide dismutase 1 (*SOD1*^*A4V/+*^) and in other fALS MNs ^3,4,7^.

To further explore cellular pathways involved in ALS pathogenesis, we employed a “patch-seq” strategy that enables neuronal excitability and gene expression to be measured simultaneously at a single cell level ^8^. Compared to prior studies where a heterogenous population of differentiated neurons were used and the neuronal maturity was not well characterized ^4,9^, we focused on a homogenous population of mature MNs that fire action potentials (APs) robustly. We introduced a GFP reporter under control of the Hb9 promoter into *SOD1*^*A4V/+*^ iPSC lines and their isogenic controls ^4^ using an AAVS1-sHb9-GFP plasmid. MNs were FACS purified after differentiation. Immunostaining for ISL1 (a MN marker) and MAP2 (a neuronal marker) revealed that most neurons had a MN identity (91.5 ± 1.4 % of 39b SOD1^A4V/+^ neurons were ISL1+ MAP2+ and 92.7 ± 1.7 % of 39b-cor SOD1^+/+^) (Extended Data Fig. 1a). To accelerate neuronal maturation, we cultured the purified MNs with glia for 3 weeks and performed whole cell current clamp recordings followed by single MN isolation for single cell RNA-seq (Smart-seq2). Motor neurons from two independent batches (Supplementary Table 1) comprising 192 neurons were analyzed, and ∼8,000 genes per neuron were detected.

There were no significant differences in housekeeping, MN, or neuronal marker gene expression between mutant and control MNs (Extended data, Fig. 1b). A significant difference in the expression of 63 genes was detected by the single cell patch-seq analysis of 39b *SOD1*^*A4V/+*^ and 39b-cor *SOD1*^*+/+*^ MNs (Fig. 1a), including downregulation of *KCNV1* (Supplementary Table 2). We confirmed this finding by a single cell patch-RT-qPCR analysis focused specifically on differential expression of 279 ion channel genes (Fig. 1b; Supplementary Table 1). There was no difference in voltage-gated Na^+^ or Ca^2+^ channel expression (Extended data, Fig. 2) and the expression signatures of most K^+^ channel subtypes were also generally very similar. However, *KCNV1*/Kv8.1 was significantly downregulated in 39b *SOD1*^*A4V/+*^ MNs relative to their isogenic controls (39b-cor *SOD1*^*+/+*^ MNs) in the single cell patch-RT-qPCR analysis, as in the patch-seq study (Fig. 1b).

**Figure 1.**
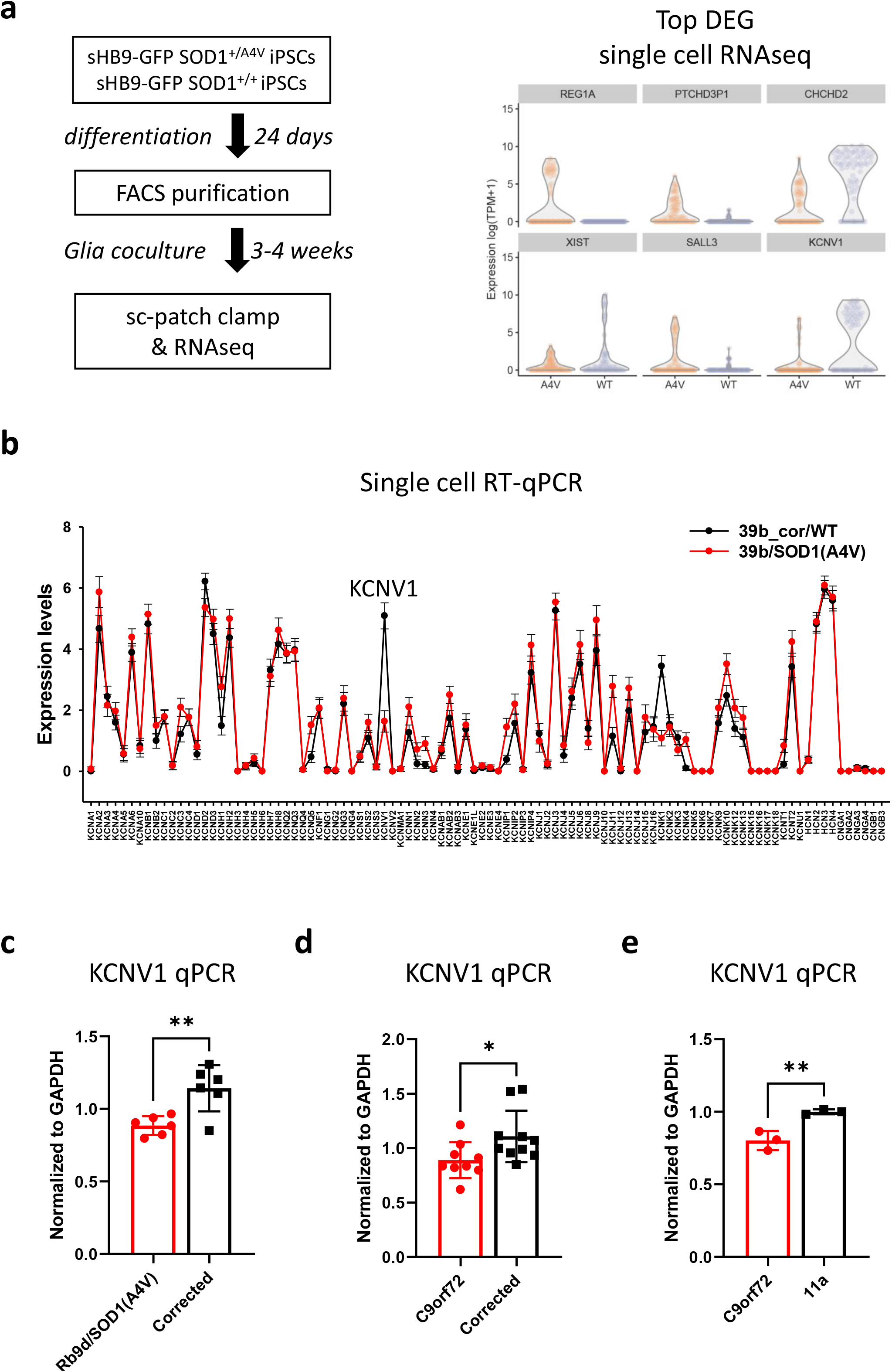
Single-cell RNA-seq analysis of 39b *SOD1*^*A4V/+*^ and 39b-cor *SOD1*^*+/+*^ MNs reveals KCNV1/Kv8.1 downregulation in mutant MNs. **a**, Single cell patch-seq scheme and top differential expressed genes identified between 39b *SOD1*^*A4V/+*^ and 39b-cor *SOD1*^*+/+*^ MNs. **b**, Comparison of K^+^ channel expression between 39b *SOD1*^*A4V/+*^ and 39b-cor *SOD1*^*+/+*^ MNs, determined by single cell multiplex RT-qPCR with ion channel probes. Average expression levels are connected to visualize differences. **c**, *KCNV1* is also downregulated in MNs from another *SOD1*^*A4V/+*^ patient’s stem cell line (RB9d *SOD1*^*A4V/+*^) compared to their isogenic control, as measured by RT-qPCR (n = 6). **d**, *KCNV1* mRNA is downregulated in C9orf72 repeat expansion patient stem cell-derived MNs compared to an isogenic control (n = 9-10). **e**, *KCNV1* mRNA is also downregulated in another C9orf72 repeat expansion patient stem cell-derived MNs compared to a healthy control line (11a) derived MNs measured by RT-qPCR (n=3). Statistical significances obtained by student’s t test (* P ≤ 0.05, ** P ≤ 0.01).

To determine whether KCNV1 downregulation is consistent across different cell lines, an independent patient derived iPSC line with the same mutation, RB9d *SOD1*^*A4V*/+^, and its isogenic control RB9d-cor *SOD1*^*+/+*^ were also investigated. RB9d *SOD1*^*A4V/+*^ MNs showed a similar downregulation of *KCNV1*/Kv8.1 compared to the control RB9d-cor *SOD1*^*+/+*^ neurons (Fig. 1c). To test if this observation expands to other ALS mutations, we examined *KCNV1* expression in MNs differentiated from iPSCs harboring C9orf72 hexanucleotide repeat expansions. We also found *KCNV1* downregulation in the C9orf72 repeat expansion iPSC MNs compared to their isogenic control MNs (provided by Dr. Coppola) (Fig. 1d), as well as in another independent C9orf72 mutant line compared to healthy controls (Fig. 1e). The downregulation of *KCNV1* in diverse ALS MN models suggests that this is not an artifact of neuronal differentiation and that it could participate in the disease progression.

*KCNV1*/Kv8.1 is a “silent” potassium channel subunit which, while it has the structure typical of voltage-activated potassium channels, cannot form functional homomeric channels. Instead, when it forms heteromeric channels with Kv2, it downregulates Kv2 channel density and modulates channel gating^8,9^. To study the role of *KCNV1*/Kv8.1 in MNs, we generated by CRISPR gene editing, a *KCNV1* knockout in control 39b-cor *SOD1*^*+/+*^ iPSCs (Extended data, Fig. 3a). Introduction of INDEL mutations resulted in early stop codon formation at the N terminal and significantly reduced *KCNV1* expression (Extended data, Fig. 3b). The *KCNV1* knockout MNs did not exhibit altered spontaneous action potential firing as assessed by multi-electrode array (MEA) recordings compared to their controls, indicating that loss of KCNV*1* does not affect neuronal excitability (Fig. 2a).

**Figure 2.**
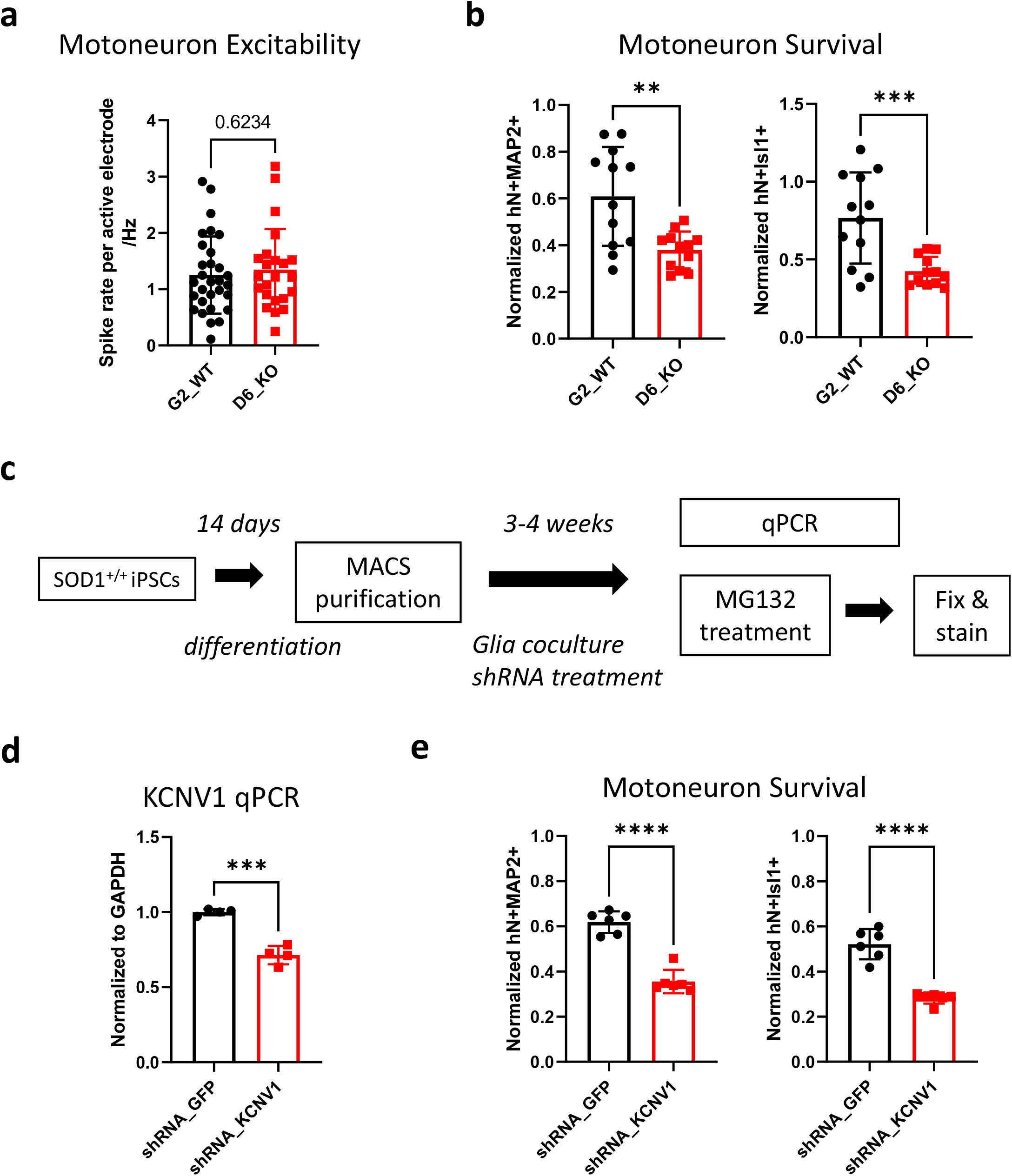
Loss of *KCNV1*/Kv8.1 increases MG132 induced neuronal cell death. **a**, Spontaneous activity of *KCNV1* CRISPR KO (D6) and control (G2) MNs recorded on a multiple electrode array (MEA). Average spike rates of active electrodes are measured (n = 23-30). **b**, After culturing MNs with glia for 4 weeks, MG-132 (10 µM) was added into the culture to induce cell death. 48h post-treatment, the cell culture was fixed and stained with MN markers: hN (human nucleus marker), MAP2 (mature neuron marker), and Isl1 (motor neuron nucleus marker). The number of hN^+^Isl1^+^ cells or hN^+^MAP2^+^ cells were quantified and normalized to DMSO treated control for each cell line. (n = 12). **c**, Outline of 2D differentiation, NCAM magnet sorting, glia coculture, shRNA treatment, followed by RT-qPCR analysis and cell death assays. **d**, *KCNV1* expression is reduced in non-mutant 39b-cor *SOD1*^*+/+*^ MNs treated with a lentivirus encoding *KCNV1* shRNA (n = 4) compared to those treated with a control GFP targeting shRNA. **e**, 39b-cor *SOD1*^*+/+*^ MNs treated with *KCNV1* shRNA show decreased survival after MG-132 treatment (n = 6). Statistical significances obtained by student’s t test (** P ≤ 0.01, *** P ≤ 0.001, **** P ≤ 0.0001).

ALS is characterized by a progressive loss of MNs, and we wondered whether *KCNV1* may impact neuronal survival. We induced cell death using MG132, a proteosome inhibitor that promotes mutant SOD1 protein aggregation ^4^. Compared to the non-gene edited wildtype controls, *KCNV1* knockout MNs exhibited a higher rate of cell death, as revealed by quantification of motor neurons post MG132 exposure (Fig. 2b). To independently confirm this result, we also knocked down *KCNV1* expression in control 39b-cor *SOD1*^*+/+*^ MNs to mimic the *KCNV1* downregulation in ALS, using two different shRNAs targeting *KCNV1*/Kv8.1, and found that knockdown of *KCNV1* in healthy MNs increased MG132 induced cell death significantly (Fig. 2c-e; Extended data Fig. 4). Downregulation of *KCNV1* in *SOD1*^*A4V/+*^ MNs, therefore increases their vulnerability to cell death without changing their excitability.

How does a reduction of *KCNV1*/Kv8.1 contribute to ALS related MN cell death? To understand this, we studied the transcriptomic alterations caused by a loss of *KCNV1*. Many genes were significantly differently expressed in *KCNV1* shRNA knockdown MNs compared to their controls (Fig. 3a; Supplementary Table 3), including genes involved in lipid metabolism, protein translation, and membrane transport pathways (Extended data Fig. 5). Gene sets related to the ER membrane, metabolism, and catabolism were among the top that were downregulated, while gene sets associated with neuron projection and intracellular transport were among the top upregulated (Fig 3b). We validated the differential expression of several of the genes (Fig. 3c), including NEK1 and OPTN, two ALS associated genes that participate in proteostasis regulation ^10,11^; RPS3A, a ribosomal and chaperone protein that counteracts α-synuclein aggregation ^12^; STMN2, a microtubule associated protein involved in TDP43 pathology ^13,14^; and VAMP3, a vesicle associated membrane protein that directs transport of proteolipid proteins ^15^. Collectively, transcriptomic analysis identified overlapping molecular pathways between ALS, and *KCNV1* knock-down MNs.

**Figure 3.**
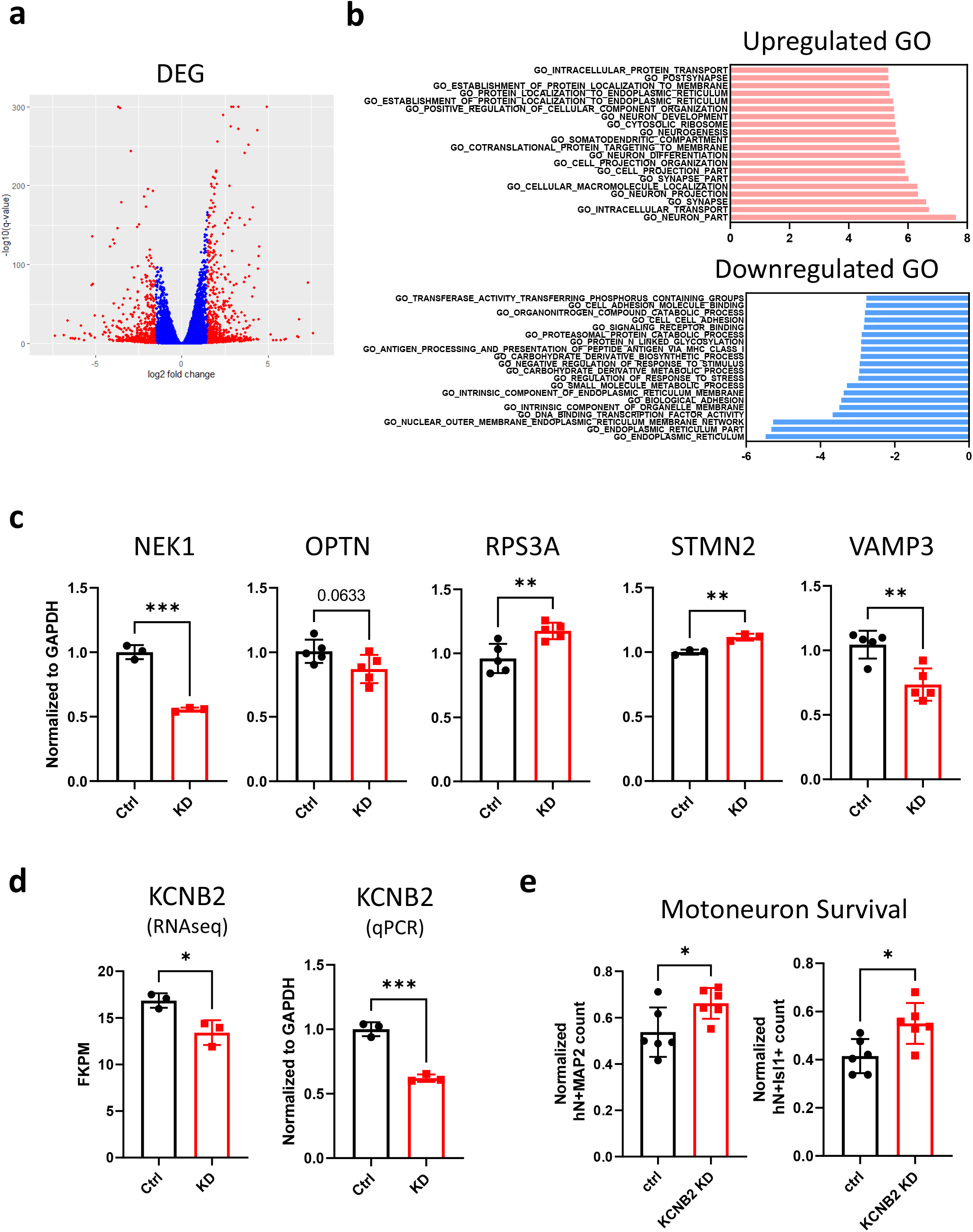
*KCNV1*/Kv8.1 interacts with the ER-PM located *KCNB2*/Kv2.2 channel and impacts multiple biological pathways. **a**, Volcano plot of differentially expressed genes (DEG) in *KCNV1* knockdown MNs. NCAM sorted MNs were treated with a lentivirus expressing *KCNV1* targeting shRNA or control *GFP* targeting shRNA and samples collected for bulk-seq analysis after 3 weeks (n = 3). The distribution between log2 (fold change KD/WT) and -log10(q-value) are plotted. Blue: log2(Fold change) < 1.5; Red: log2(Fold change) ≥ 1.5. **b**, Top enriched gene sets in *KCNV1* knock down MNs. Red: upregulated; Blue: downregulated. X-axis: normalized enrichment score. c, RT-qPCR validation in *KCNV1* knockdown MNs (n = 3-5). **d**, Expression of the KCNB2/Kv2.2 transcript detected by RNAseq (left, n = 3) and RT-qPCR (right, n = 3). **e**, 39b *SOD1*^*A4V/+*^ MNs treated with KCNB2 shRNA show increased survival after MG-132 treatment (n = 6). Statistical significance obtained by student’s t test (* P ≤ 0.05, ** P ≤ 0.01, *** P ≤ 0.001, **** P ≤ 0.0001).

KCNV1/Kv8.1 forms heteromers with Kv2 channels, and we therefore examined if Kv2 was involved in the MN cell death induced by a reduction of KCNV1 expression. While channels formed by Kv2 subunits regulate neuronal excitability ^16,17^, they also have non-ion channel functions, including chaperoning proteins to the plasma membrane ^18^ and coupling the plasma membrane (PM) to the endoplasmic reticulum (ER) membrane^19,20^ as well as producing pro-apoptotic effects under conditions of oxidative stress ^21-23^. Recent studies show that ER-PM located Kv2 ion channels regulate ER Ca^2+^ uptake and release in neurons ^24-26^ as well as lipid metabolism ^27^ which could impact multiple biological processes, including synaptic transmission, receptor signaling, membrane trafficking, and cytoskeletal dynamics ^28^. We found that the expression of *KCNB2* which encodes the Kv2.2 ion channel was significantly decreased in *KCNV1* knock down MNs (Fig. 3d). We wondered whether this was protective or pathogenic. To explore this, we knocked down *KCNB2*/Kv2.2 expression in 39b *SOD1*^*A4V/+*^ALS MNs, and then induced cell death using MG-132. We found that compared to controls, survival of ALS MNS was significantly increased by KNCB2 shRNA knockdown (Fig. 3e). Suppression of KCNB2 was recently reported to rescue the eye degeneration found in a C9orf72 expansion drosophila model ^29^. It is possible, therefore, that a loss of KCNV1/Kv8.1 increases MN vulnerability through an interaction with Kv2.2 ion channels, and that the transcriptional downregulation of Kv2.2 in KCNV1/Kv8.1 knockdown MNs is a protective compensatory response.

Potassium ion channels are the most diverse and widely distributed ion channels in mammals and play critical roles in controlling membrane excitability. Recently, more diverse functions of potassium channels have been reported, suggesting they also modulate processes such as cellular Ca^2+^ metabolism ^24,25^, lipid metabolism ^27^, apoptosis ^30^, and neuron-microglia interactions ^31^. Our data now reveal that a loss of the silent subunit Kv8.1 increases ALS MN vulnerability. We suggest this may occur through an interaction with the ER-PM located Kv2 channel that leads to the dysregulation of multiple pathways, including lipid metabolism and membrane transport. The precise underlying molecular machinery of Kv8.1-Kv2 interactions in ALS MNs now needs to be defined, and this may open opportunities for novel therapeutic interventions.

## Supporting information

Supplemental Tabls

## Acknowledgments

We acknowledge Target ALS, ALS Alliance, an HBI ALSA grant, and DoD Grant W81XWH2010077 for support of this study. We acknowledge Dr. Giovanni Coppola for kindly providing the C9orf72 expansion iPSC line as well as the corrected isogenic control line.

## Conflict of Interest

CJW and KE are founders of QurAlis Corporation.

Supplementary Table 1: Experimental design of single cell patch-seq and patch-qPCR study

Supplementary Table 2: List of 63 DEG genes identified from single cell patch-seq study in 39b *SOD1*^*A4V/+*^ and 39b-cor *SOD1*^*+/+*^ MNs

Supplementary Table 3: DEG list of bulk-seq collected from 39b-cor *SOD1*^*+/+*^ MNs treated with *KCNV1* and control shRNAs

**Extended Data Figure 1.**
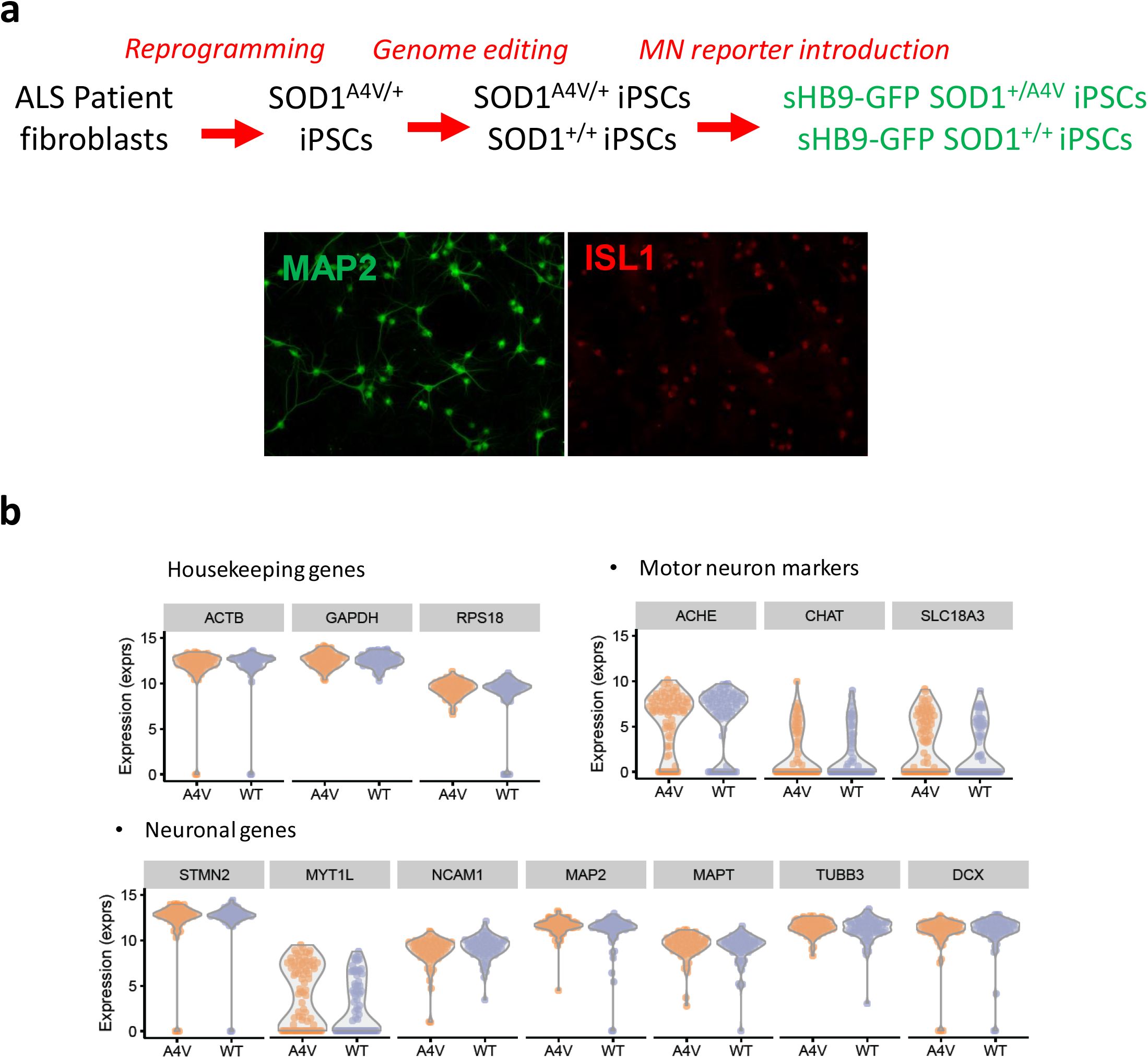
Generation of 39b *SOD1*^*A4V/+*^ and 39b-cor *SOD1*^*+/+*^ reporter iPS cell lines and their differentiation into motor neurons (MN) for single cell profiling. **a**, Schematic representation of *SOD1*^*A4V/+*^ ALS patient iPSC generation, isogenic correction using genome editing, and introduction of the Hb9 MN reporter. Hb9-GFP positive MNs maintain their identity as indicated by ISL1 expression when co-cultured with mouse primary glial cells for 24 days. **b**, After recording and picking of MNs, cDNA libraries were constructed and sequenced. Violin plot shows that expression of housekeeping genes, motor neuron marker, and neuronal genes is similar between genotypes.

**Extended Data Figure 2.**
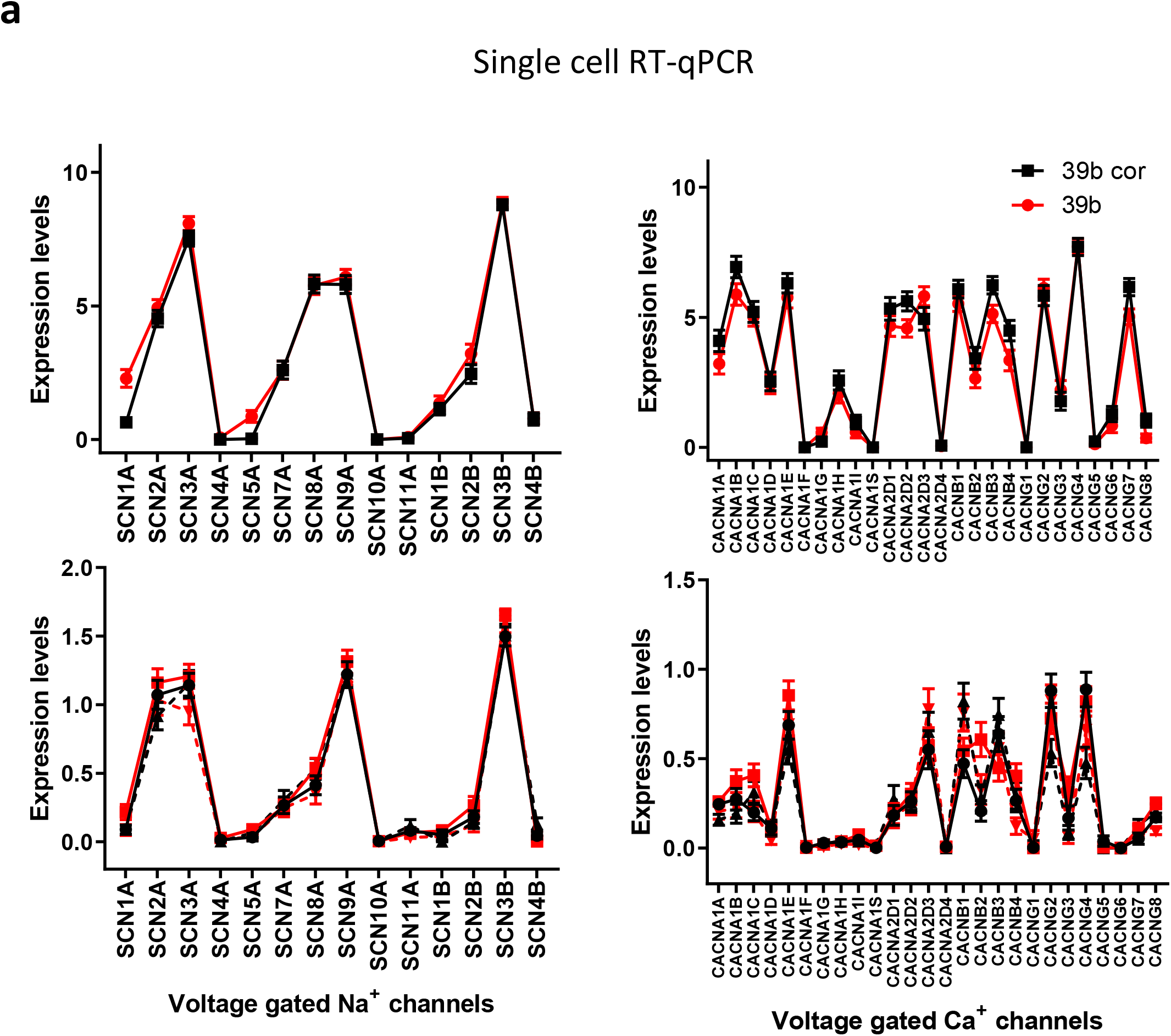
**a**, Comparison of Na^+^ and Ca^2+^ channel expression between 39b SOD1^A4V/+^ and 39b-cor SOD1^+/+^ MNs determined by single cell RT-PCR.

**Extended Data Figure 3.**
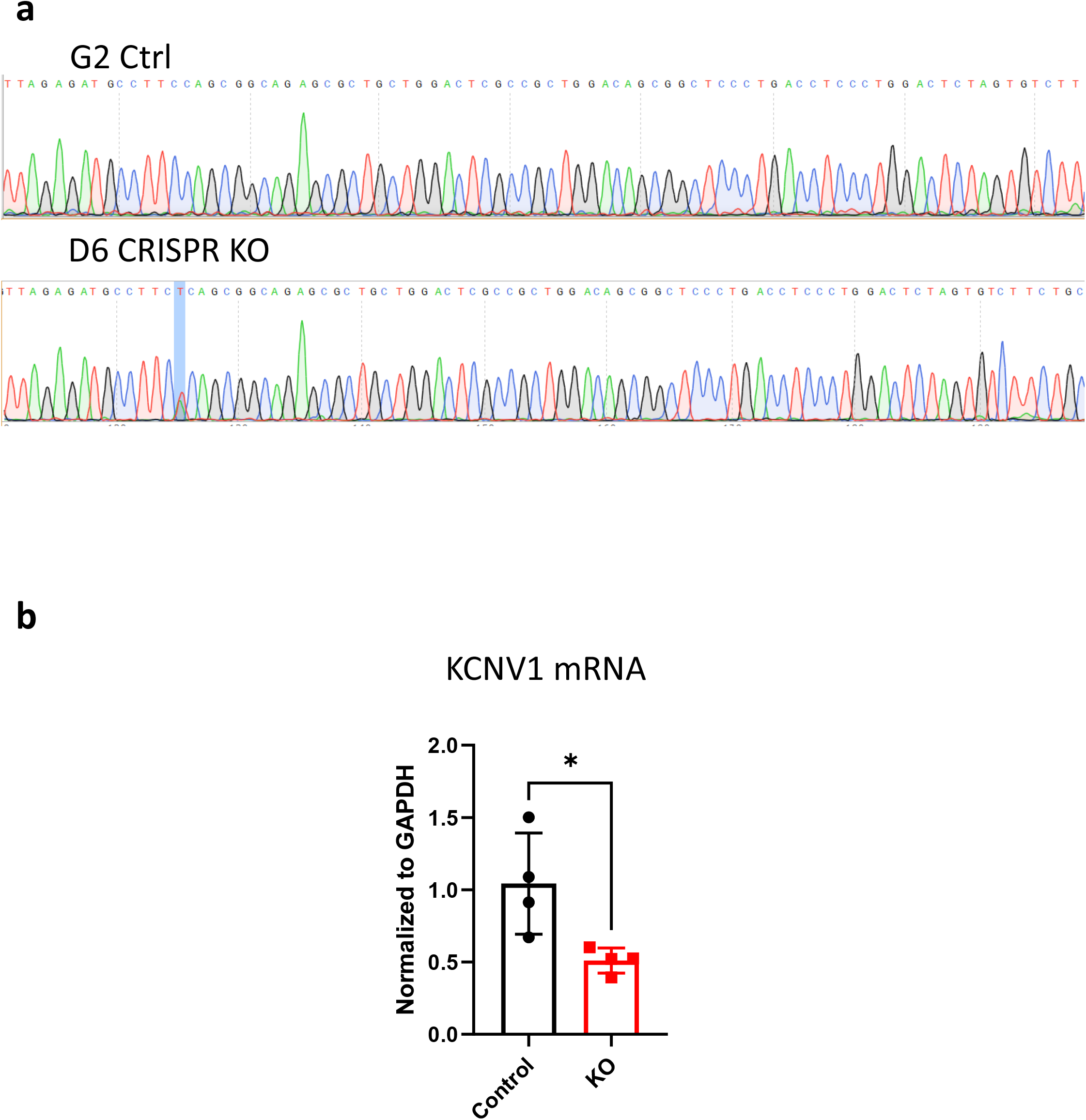
Generation of the KCNV1 CRISPR KO. **a**, Sequence of *KCNV1* gene in G2 (control) and D6 (KO) iPSC lines. Single nucleotide insertions were generated in D6 iPSC. **b**, The expression of *KCNV1* mRNA quantified by qRT-PCR in MNs differentiated from G2 and D6 iPSC (n=4).

**Extended Data Figure 4.**
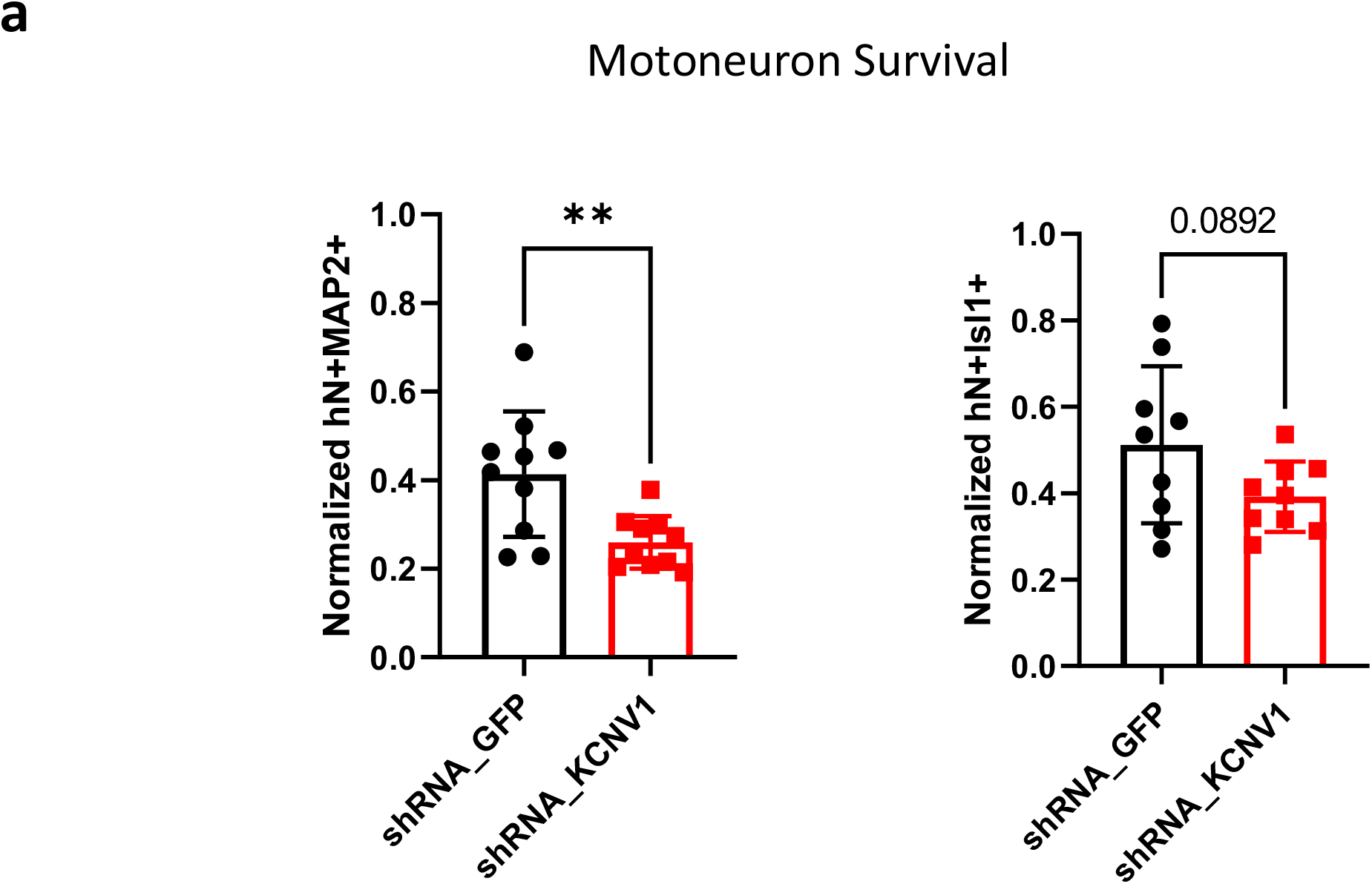
**a**, 39b-cor *SOD1*^*+/+*^ MNs treated with another *KCNV1* shRNA also show decreased survival after MG-132 treatment (n = 9-10). Statistical significances obtained by student’s t test (** P ≤ 0.01).

**Extended Data Figure 5.**
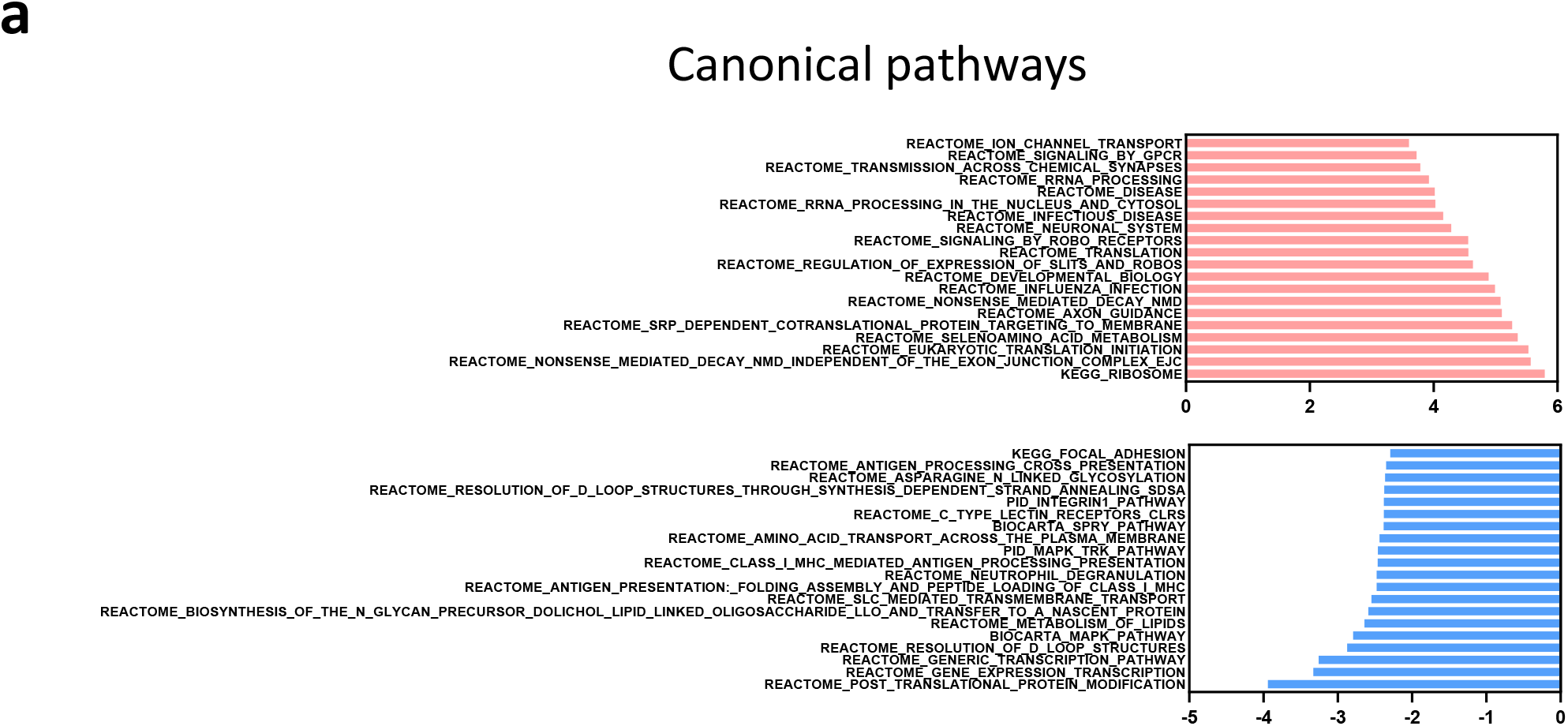
**a**, Top enriched pathways in *KCNV1* knock down MNs. Red: up-regulated; Blue: down-regulated. X-axis: normalized enrichment score

## Materials and methods

### iPSC cultures

The following clonal iPSC lines were used in this study: patient iPSCs (39b SOD1^A4V/+^, Rb9d SOD1^A4V/+^, C9ORF72 expansion), their isogenic controls iPSCs (39b-cor SOD1^+/+^, Rb9d-cor SOD1^+/+^, C9ORF72 corrected), and a healthy wildtype line 11a. iPSCs were maintained with mTeSR1 media (Stem Cell Tech) or StemFlex media (Gibco #A3349401) on culture dishes coated with Matrigel (BD Biosciences). Media was changed daily, and cells were typically passaged by accutase. All cell cultures were maintained at 37°, 5% CO_2_.

### HB9::GFP transfection using zinc finger nuclease (ZFN)

For transfection of a HB9::GFP reporter into each of the four iPSC lines, a 1kb HB9 promoter fragment (gift from Hynek Wichterle) controlling the expression of myristoylated GFP was inserted into a donor plasmid specific for the AAVS1 locus (Sigma). Subsequently, 2.5 million iPSCs were treated with accutase and electroporated using the Neon transfection system (100 μl tip; 1600 V Voltage, 20 ms Width, 1 Pulse; Life Technologies) with 2 μg of AAVS1 ZFN plasmid and 6 μg of the modified AAVS1 donor plasmid. After nucleofection, cells were plated on matrigel with mTeSR1 in the presence of 10 μM ROCK inhibitor. After 48hrs puromycin selection, surviving clonal colonies were individually passaged and gDNA was extracted. PCR was used to confirm proper targeting of the cassette. Expression of the reporter was verified using expression of GFP and the motor neuron marker Isl1.

### Motor neuron differentiation

Motor neuron differentiation for patch-seq and patch-qPCR was carried out as described previously ^32^ with modifications ^33^ in a 24-day protocol based on initial neuralization with SB431542 (10 μM, Sigma Aldrich) and Dorsomorphin (1 μM, Stemgent), and motor neuron patterning with retinoic acid (1 μM, Sigma) and a small smoothened Agonist 1.3 (1 μM, Calbiochem). Differentiated motor neurons were dissociated using accutase, then filtered with a 70 µm filter and motor neurons, subsequently, isolated by activated flow cell sorting using HB9::GFP expression. Motor neurons were plated in the presence of P0 mouse glial cells and were maintained in Neurobasal media, supplemented with N2 and B27 (Invitrogen), 10 ng/mL each of BDNF, GDNF, CNTF (R&D) and ascorbic acid (0.4 μg/ml, Sigma) and fed every 2-3 days. Motor neuron differentiation for cell death assays and RT-qPCR was carried out with a modified 14-day protocol described in Klim et al ^14^. Briefly, iPS cells were dissociated using accutase and plated with 1.5 million cells per 10 cm dishes coated with geltrex. 1 day after plating, the medium was changed to differentiation medium (half Neurobasal and half DMEM/F12 supplemented with B27 and N2 supplements, GlutaMax, and non-essential amino-acids) from StemFlex™ Medium (Gibco). 10 µM SB431542, 100 nM LDN-193189, 1 µM retinoic acid and 1 µM Smoothend agonist were treated on d0-5, and 5 µM DAPT, 4 µM SU-5402, 1 µM retinoic acid, and 1 µM Smoothend agonist were treated on d6-14. The differentiated cells were dissociated using accutase, then filtered with a 70 µM filter. The single cell suspension was incubated with PE Mouse Anti-Human CD56 (BD Biosciences #555516) and then Anti-R-Phycoerythrin (PE) Magnetic Particles (BD Biosciences #557899) in MACS buffer (PBS with 2mM EDTA and 0.5% BSA) on ice. The sorted motor neurons were plated in the presence of P0 mouse glial cells and were maintained in Neurobasal media, supplemented with N2 and B27 (Invitrogen), 10 ng/mL each of BDNF, GDNF, CNTF (R&D) and ascorbic acid (0.4 μg/ml, Sigma) and fed every 2-3 days.

### Immunocytochemistry

Cell cultures were fixed in 4% PFA for 15minutes at 4°C, permeabilized with 0.2% Triton-X in PBS for 45 minutes and blocked with 10% donkey serum in PBS-T (Triton 0.1%). Cells were then incubated in primary antibody overnight and secondary antibodies for 1 hour in 2% donkey serum in PBS-T after several washes in between. The following antibodies were used: primary antibodies, human nuclei (1:1000, Millipore), Islet1 (1:200, DSHB, 40.2D6), and MAP2 (1:10000, Abcam ab5392), and secondary antibodies AlexaFluor (1:1000, Life Technologies) and DyLight (1:500, Jackson ImmunoResearch Laboratories). Images were acquired and analyzed by the ArrayscanTM XTI (Thermo Fisher). To induce cell death, motor neuron-mouse glia coculture were maintained for 3-4 weeks, then treated with 10 µM MG132 for 48 hours before fixation and staining.

### Electrophysiology

Current clamp on iPSC-derived MNs: The purified and differentiated neurons on P0 mouse glial cells were identified microscopically and whole cell current clamp recordings were conducted using a Multiclamp 700B amplifier. Data were digitized with a Digidata 1440A A/D interface and recorded using pCLAMP10 (Molecular Devices). Borosilicate glass pipettes were pulled on a P-97 puller (Sutter Instruments). The external solution consisted of 140 mM NaCl, 5 mM KCl, 2 mM CaCl_2_, 2 mM MgCl_2_, 10 mM HEPES, and 10 mM D-glucose, pH 7.4, adjusted with NaOH. The internal solution consisted of 135 mM K-Gluconate, 10 mM KCl, 1 mM MgCl_2_, 5 mM EGTA, and 10 mM HEPES. Fast and slow capacitance transients and whole cell capacitance were compensated using the automatic capacitance compensation in voltage clamp mode on the Multiclamp 700B. After currents were injected to bring the membrane potential to -60 mV in current clamp mode, a depolarizing current ramp (0 to 700 pA in one second) or series of steps (500-ms steps from 0 to 100 pA) were applied to measure the firing frequency and the action potential characteristics. Spikes were counted using a criterion of a peak voltage > -10 mV and amplitude > 20mV. Frequency of firing was calculated from all the spikes during stimulation and instant frequency from the first two spikes. Action potential characteristics including trough voltage were analyzed from the first action potential waveform.

### Single cell collection

For patch-qPCR, single neurons were picked immediately after recording, transferred into 8 well strip tubes containing 5 μl lysis buffer (CellsDirect™ One-Step qRT-PCR Kit, #11753100), then placed on dry ice. Tubes were kept at –80 °C until the next pre-amplification step. For patch-seq, the single neurons were collected in 5μl of 1x TCL buffer (Qiagen, #1031576) in each well of 96 well PCR plates (Eppendorf, #951020401) on dry ice and then kept at -80 °C until complementary DNA (cDNA) construction.

### Single cell RNA-seq

Single cell RNA sequencing libraries were generated using the SmartSeq2 protocol^34^ with minor modifications ^35^. All libraries were prepared by the Broad Technology Labs and sequenced at the Broad Genomics Platform. Briefly, total RNA from single cells was purified using RNA-SPRI beads. Poly(A)+ mRNA was converted to cDNA and amplified. cDNA was subjected to transposon-based fragmentation using dual-indexing to barcode each transcript fragment with a combination of barcodes specific to a single cell. Barcoded cDNA fragments were then pooled for sequencing. Sequencing was carried out as paired-end 2×25bp with an additional 8 cycles for each index. To obtain expression values for each cell, the data was demultiplexed and aligned to the human genome (hg19) using Tophat version 2.0.10 with default settings ^36^. Transcripts were quantified by the BTL computational pipeline using Cuffquant version 2.2.1 ^37^ and raw counts of reads mapped per gene were extracted from aligned bam using Feature Counts (subread-1.5.0-p1) with default settings. For visualization purposes expression levels were converted to log-space by taking the log2(FPKM+1) or by normalizing read counts to transcripts per million using log2 (TPM+1). To identify genes that are differentially expressed between distinct electrophysiological classes or genotypes, we first filtered cells with poor electrophysiological recordings or low sequencing quality (library size greater or less than 3 median-absolute-deviations from the median). Single cell differential expression analysis was performed using SCDE ^38^ with default settings and batch effects were corrected for when calculating expression differences.

### Single-cell RT-qPCR: Primer design

DELTAgene assays (Fluidigm) were designed for 279 genes (listed in the Supplement Table 3), including housekeeping genes (GAPDH and ACTB), motor neuron markers (MNX1, CHAT, ISL1, and SLC18A3), a glial cell marker (mGfap), voltage-gated ion channels, and ligand-gated ion channels. The assays are designed to cross an intron. A nested primer strategy (outer and inner pairs of forward and reverse primers) was utilized. Outer primer pairs were used for the pre-amplification and inner primer pairs were used for qRT-PCR in the Biomark. For outer primers, the oligos were synthesized by IDT, dissolved at a concentration of 200 μM in H_2_O, and 267 gene primer pairs mixed in a 15 ml falcon tube at a final concentration of 200 nM each primer. For inner primers, the oligos were synthesized by IDT and dissolved at a concentration of 50 μM in H_2_O. To make 10 μl of 20 μM primer pairs per each gene in V-shaped 96 well plate (Eppendorf, #951020401), 4 μl of forward and reverse primers were added to 2 μl H_2_O.

### Single-cell RT-qPCR: cDNA synthesis and Pre-amplification

We used a protocol adapted from Fluidigm (Application Note MRKT00075e) that combines reverse transcription and preamplification (called RT-STA, reverse transcription-specific target amplification) using CellsDirect™ One-Step qRT-PCR Kit. A mixture consisting of 2.8 μl of multiplex outer primers plus 0.2 μl of the enzyme mix (SuperScript® III Reverse Transcriptase and Platinum® Taq DNA Polymerase) was added to each single cell in lysis buffer in the 8 well strips. Following a centrifugation, the 8 well strips were moved to a thermal cycler and subjected to the following protocol: 50 °C, 60 min for reverse transcription; 95 °C, 2 min for Taq enzyme activation; 20 cycles of (95 °C, 15 sec; 60 °C, 4 min). The reactions were kept at –80 °C until the next tests.

To determine a proper dilution for following single cell qPCR, a few randomly picked reactions were diluted in H_2_O at 25-, 50-, 75- and 100-folds and tested with GAPDH, a housekeeping gene, by qPCR. A dilution of 25-fold was selected for subsequent experiments. All the reactions were diluted and tested with GAPDH, a positive control, and mGfap, a negative control, to determine whether we had properly picked neurons without glial cells.

### Single-cell RT-qPCR: Data acquisition and analysis

Diluted reactions were analyzed by qPCR using 96.96 Dynamic ArrayTM Integrated Fluidic Circuits (IFCs) and the Biomark™ HD system from Fluidigm. The IFCs were processed and the instruments were operated in the BCH IDDRC Molecular Genetics Core Facility according to the manufacturer’s procedures for analyzing DELTAgene assays. Three IFCs were used to analyze the 96 samples for the total of 267 assays, and all IFCs included housekeeping genes GAPDH and ACTB, and a blank. In single-cell qPCR analysis, a Ct of 30 was used as the background value for all real-time signals. Expression levels were calculated by 30 – Ct value, assigning a value of 0 when Ct > 30. Hierarchical clustering and heatmap were generated with R software.

### CRISPR KO generations for KCNV1/Kv8.1

39b-cor SOD1^+/+^ iPSCs were maintained in geltrex-coated 10 cm dishes in StemFlex™ Medium (Gibco, #A3349401) at 37⍰°C and 5% CO^2^. Two sgRNA CUGGACUCGCCGCUGGACAG and GGGUUAGAGAUGCCUUCCAG were designed and synthesized from Synthego to create INDEL in KCNV1. 39b-cor SOD1^+/+^ iPSC were dissociated into single-cell suspension using⍰accutase with 10 µM Y-27632. 1 million cells were nucleofected with 1.2 µl of 20 µM Cas9-2NLS and 9 µl of 30 µM each sgRNA using Nucleofactor™ device (program CB1150, Lonza) according to the manufacturer’s instructions. Transfected cells were plated on geltrex-coated 6 well plates in StemFlex™ Medium with 10 µM Y-27632. After two days, the cells were dissociated into single cells and plated at a density of 2000 cells/10⍰cm dish. 1 week after transfection, single colonies were picked and sequenced to confirm INDEL. Forward and reverse primers were 5’TGCGCCAAGGAGAGGTAA3’ and 5’TCGCTGCAGAAGACACTAGA3’, respectively.

### RNA extraction and RT-qPCR

RNA was extracted from motoneuron culture using RNeasy Micro Kit (Qiagen) and then cDNA was generated using Superscript Vilo Synthesis Kit (Invitrogen) following manufacturer’s manuals. qPCR was performed on Applied Biosystems 7500 machine (Life Technologies) using Fast SYBR Green Master Mix (Roche).

### RNAseq analysis

In this study, RNA-sequencing libraries were prepared using the TruSeq Stranded RNA kit with RiboZero Gold to enrich for messenger RNA (mRNA). The libraries were then sequenced on a NovaSeq6000 platform, generating paired-end reads of 100 base pairs in length. To ensure the accuracy of downstream analysis, the reads were classified into five categories using Xenome (v1.0.0), a tool designed to remove reads derived from unwanted sources such as contaminating DNA or RNA from other species. Reads classified as human were then aligned to the human genome (Hg38) using STAR aligner (v2.7.5c). Average input read counts were 31.00±3.61M(SD) and average percentage of uniquely aligned reads were 88.04±0.03(SD)%. Total counts of read-fragments aligned to known gene regions within the human ensemble gene model annotation (GRCh38) are used as the basis for quantification of gene expression. Fragment counts were derived using HTSeq program (ver 0.12.4). Quality control measures were performed to assess the quality of the data, including base quality, mismatch rate, and mapping rate to the whole genome. Additionally, repeats, chromosomes, key transcriptomic regions (exons, introns, UTRs, genes), insert sizes, AT/GC dropout, transcript coverage, and GC bias were assessed to identify potential issues in the library preparation or sequencing. To identify genes that were differentially expressed between conditions, lowly expressed genes were removed and genes with counts per million (CPM) greater than 0.15 in at least three samples were selected for downstream analysis. Differential expression analysis was conducted using the Bioconductor package EdgeR, (ver 3.14.0) which uses a negative binomial model to account for the variability in the data. Expressed genes were sorted by directional pVal and used as input for Gene set enrichment analysis (GSEA). MsigDB (ver7.0) was used as reference gene set. Raw and processed data were deposited within the Gene Expression Omnibus (GEO) repository (www.ncbi.nlm.nih.gov/geo).

